# Stability of DNA methylation and chromatin accessibility in structurally diverse maize genomes

**DOI:** 10.1101/2021.03.10.434810

**Authors:** Jaclyn M Noshay, Zhikai Liang, Peng Zhou, Peter A Crisp, Alexandre P Marand, Candice N Hirsch, Robert J Schmitz, Nathan M Springer

**Author notes:** Corresponding author: Nathan Springer.

## Abstract

Accessible chromatin and unmethylated DNA are associated with many genes and cis-regulatory elements. Attempts to understand natural variation for accessible chromatin regions (ACRs) and unmethylated regions (UMRs) often rely upon alignments to a single reference genome. This limits the ability to assess regions that are absent in the reference genome assembly and monitor how nearby structural variants influence variation in chromatin state. In this study, *de novo* genome assemblies for four maize inbreds (B73, Mo17, Oh43 and W22) are utilized to assess chromatin accessibility and DNA methylation patterns in a pan-genome context. The number of UMRs and ACRs that can be identified is more accurate when chromatin data is aligned to the matched genome rather than a single reference genome. While there are UMRs and ACRs present within genomic regions that are not shared between genotypes, these features are substantially enriched within shared regions, as determined by chromosomal alignments. Characterization of UMRs present within shared genomic regions reveals that most UMRs maintain the unmethylated state in other genotypes with only a small number being polymorphic between genotypes. However, the majority of UMRs between genotypes only exhibit partial overlaps suggesting that the boundaries between methylated and unmethylated DNA are dynamic. This instability is not solely due to sequence variation as these partially overlapping UMRs are frequently found within genomic regions that lack sequence variation. The ability to compare chromatin properties among individuals with structural variation enables pan-epigenome analyses to study the sources of variation for accessible chromatin and unmethylated DNA.

**Article summary:** Regions of the genome that have accessible chromatin or unmethylated DNA are often associated with cis-regulatory elements. We assessed chromatin accessibility and DNA methylation in four structurally diverse maize genomes. There are accessible or unmethylated regions within the non-shared portions of the genomes but these features are depleted within these regions. Evaluating the dynamics of methylation and accessibility between genotypes reveals conservation of features, albeit with variable boundaries suggesting some instability of the precise edges of unmethylated regions.

## Introduction

The 2.1Gb maize B73 genome was first assembled in 2009 and contains ~80% repetitive sequence (Schnable *et al.* 2009). Unlike model species such as *Arabidopsis thaliana*, maize has transposable elements and highly methylated regions that are interspersed with genic regions of the genome (The Arabidopsis Genome Initiative 2000; Baucom *et al.* 2009; Springer and Schmitz 2017). One challenge in complex crop genomes such as maize is the identification of regulatory elements within genomes. There are opportunities to utilize both chromatin properties such as DNA methylation or chromatin accessibility to identify functional elements.

The maize genome is highly methylated and regions containing DNA methylation can be sub-classified based on the specific sequence context of the methylation. High levels of CG and CHG (H = A, C or T) methylation without CHH methylation are often found over transposable elements and other repetitive regions of the maize genome, while CG-only methylation is observed frequently within gene bodies (West *et al.* 2014; Niederhuth *et al.* 2016; Crisp *et al.* 2020). CHH methylation, which is largely the result of RNA-directed DNA methylation (RdDM), is found near highly expressed genes (Gent *et al.* 2013; Li *et al.* 2015a; Niederhuth *et al.* 2016). A small proportion of the maize genome lacks DNA methylation in any sequence context and these unmethylated regions (UMRs) likely reflect regions with potential roles in regulation of gene expression (Oka *et al.* 2017; Ricci *et al.* 2019; Hoefsloot and Stam 2020; Crisp *et al.* 2020).

Chromatin accessibility is another feature of chromatin that can be used to identify genomic regions with roles in regulation of transcription. In maize, ~1% of the genome contains accessible chromatin when profiled with a single tissue type (Rodgers-Melnick *et al.* 2016). Profiles of chromatin accessibility combined with other chromatin modifications have identified potential regulatory elements in the maize genome (Oka *et al.* 2017; Ricci *et al.* 2019). While chromatin accessibility is quite useful for identifying regulatory elements in a particular tissue, this property is highly dynamic with changes between tissue types or cells (Ricci *et al.* 2019; Marand *et al.* 2020a; Crisp *et al.* 2020). The vast majority of accessible chromatin occurs in unmethylated regions of the genome. However, there are additional unmethylated regions that do not exhibit chromatin accessibility. These likely reflect the fact that the unmethylated regions of the genome are quite stable in vegetative tissues while chromatin accessibility is highly tissue-specific (Schmitz *et al.* 2013; Kawakatsu *et al.* 2016; Marand *et al.* 2020b; Crisp *et al.* 2020). To date, the analysis of chromatin accessibility in maize has largely focused on the accessible regions within the B73 genome.

The analysis of chromatin properties within the B73 reference genome has been useful for functional annotation of the genome. However, there is also value in assessing natural variation for the chromatin properties in different inbred lines of maize. While chromatin accessibility studies have largely focused on B73, many studies have compared DNA methylation between maize genotypes (Eichten *et al.* 2013; Regulski *et al.* 2013; Li *et al.* 2015b; Anderson *et al.* 2018; Xu *et al.* 2019, 2020). These studies have found many examples of DNA methylation variation. Changes in DNA methylation can occur due to alterations in DNA sequence such as transposon insertions (Noshay *et al.* 2019) or can occur in regions with no genetic changes (Eichten *et al.* 2011). The ability to fully compare DNA methylation patterns among genotypes and to investigate the role of structural variation has been limited due to reliance upon a single reference genome for comparisons.

The genome content varies substantially among maize genotypes (Fu and Dooner 2002; Springer *et al.* 2009; Swanson-Wagner *et al.* 2010; Anderson *et al.* 2019; Hufford *et al.* 2021). The availability of multiple *de nov*o assembled reference genomes has enabled whole genome comparisons of genome content (Hirsch *et al.* 2016; Springer *et al.* 2018; Sun *et al.* 2018; Haberer *et al.* 2020; Hufford *et al.* 2021). Many of the sequences present in any one inbred are not present at collinear regions in other genomes (Fu and Dooner 2002; Sun *et al.* 2018; Haberer *et al.* 2020). This results in a pan-genome that contains more genes and transposons than any individual maize inbred (Hirsch *et al.* 2014; Anderson *et al.* 2019; Hufford *et al.* 2021). While it is quite clear that genome content differs substantially, it has been difficult to assess the chromatin of the pan-genome due to technical difficulties in connecting the same sequence regions between genotypes.

In this study, we generated DNA methylation and chromatin accessibility profiles from four maize inbreds that each have *de novo* genome assemblies. UMRs and ACRs are identified for each genotype based on alignment of the chromatin data to the B73v4 genome and the genome from which it was generated. Chromosomal alignments were used to classify shared and nonshared sequences between genomes. UMRs and ACRs are substantially depleted within the non-shared portions of the genome. We assessed the stability of UMRs between genotypes within the shared regions of the genome. While the majority of UMRs in these regions have an overlapping UMR in another genotype, the majority do not have identical boundaries. These UMRs with shifted boundaries account for a large portion of the differentially methylation regions between two genotypes. The partially overlapping UMRs are not enriched for variable chromatin accessibility or changes in expression of nearby genes, suggesting that differences in the specific boundaries between methylated and unmethylated DNA are tolerated with little functional impact.

## Results

### Characterization of unmethylated DNA and accessible chromatin in four maize genomes

DNA methylation (profiled using whole genome bisulfite sequencing - WGBS (Cokus *et al.* 2008; Lister and Ecker 2009)), chromatin accessibility (profiled using Assay for Transposase Accessible Chromatin-sequencing - ATAC-seq (Buenrostro *et al.* 2013)), and gene expression (RNA-seq) data were generated for the same tissue sample from seedling leaf of four maize inbreds (B73, Mo17, W22 and Oh43) (Table S1). For all genotypes the resulting datasets were aligned to their own genome assembly and non-B73 genotypes were additionally aligned to the B73v4 reference genome assembly.

The alignment rates for the WGBS datasets were substantially higher when mapped to their respective genome assembly (~60%) compared to non-B73 samples mapped to the B73 reference genome assembly (~43%) (Table S1). The reduced mapping rate when aligning data from non-B73 genotypes to the B73 genome assembly is likely due to polymorphisms and structural variants present between inbreds. We focused on analysis of methylation classifications based analysis of merged replicates, since the data from the two biological replicates was highly correlated and the unmethylated regions identified within individual samples were frequently (>97%) found in the merged sample (Table S3). The WGBS data was used to classify the methylation state for each 100bp bin based on context-specific DNA methylation (Figure S1A) as described previously (Crisp *et al.* 2020). Bins were classified as unmethylated (<20% methylation in all contexts), CHH (CHH>15%), CG/CHG (>40% both CG and CHG), CG only (>40% CG), missing data, missing sites or intermediate methylation (Figure S1A). The majority (71-74%) of the maize genome is classified as methylated with most of this exhibiting CG/CHG methylation and in rare cases CHH methylation (Figure S1A). A much smaller proportion (6-7%) of the genome is classified as unmethylated (Figure S1A). In each genome, roughly 15% of the bins are classified as missing data, likely due to an inability to align WGBS reads uniquely to repetitive regions. However, the proportion of bins with missing data was substantially larger when non-B73 WGBS data was aligned to the B73 genome (Figure S1B).

The unmethylated 100bp bins were merged and filtered (Crisp *et al.* 2020) to identify unmethylated regions (UMRs) (Table 1). UMRs were defined for each inbred based on alignment to their respective genome assembly and when aligned to B73 (Figure 1A). The total number of UMRs was similar across all four genotypes, although a greater number of UMRs were defined when mapping WGBS reads to the sample-matched genome assembly. UMRs were classified as genic, proximal (<2kb from nearest gene) and intergenic (>2kb from nearest gene) in all four genotypes based on alignment of samples to their cognate reference genome assembly, and were consistent across genotypes with ~50% of UMRs being observed in genic regions and ~40% in intergenic regions (Figure 1B).

**Table 1:** UMR and ACR summary statistics

| Sample Genotype | Ref Genotype | # of bins defined as Missing Data | # of bins defined as Methylated* | # of bins defined as Unmethylated | # of UMRs | # of ACRs |
| --- | --- | --- | --- | --- | --- | --- |
| B73 | B73v4 | 3511785 | 15064391 | 1325187 | 107178 | 24304 |
| Mo17 | Mo17 | 3649729 | 15566698 | 1385916 | 113838 | 24309 |
| Oh43 | Oh43 | 3096596 | 15719767 | 1445686 | 111261 | 22774 |
| W22 | W22 | 3322802 | 15315985 | 1369207 | 112253 | 21232 |

\*Methylated is the combined value of bins defined as CG only, CG/CHG, and CHH

**Figure 1:**
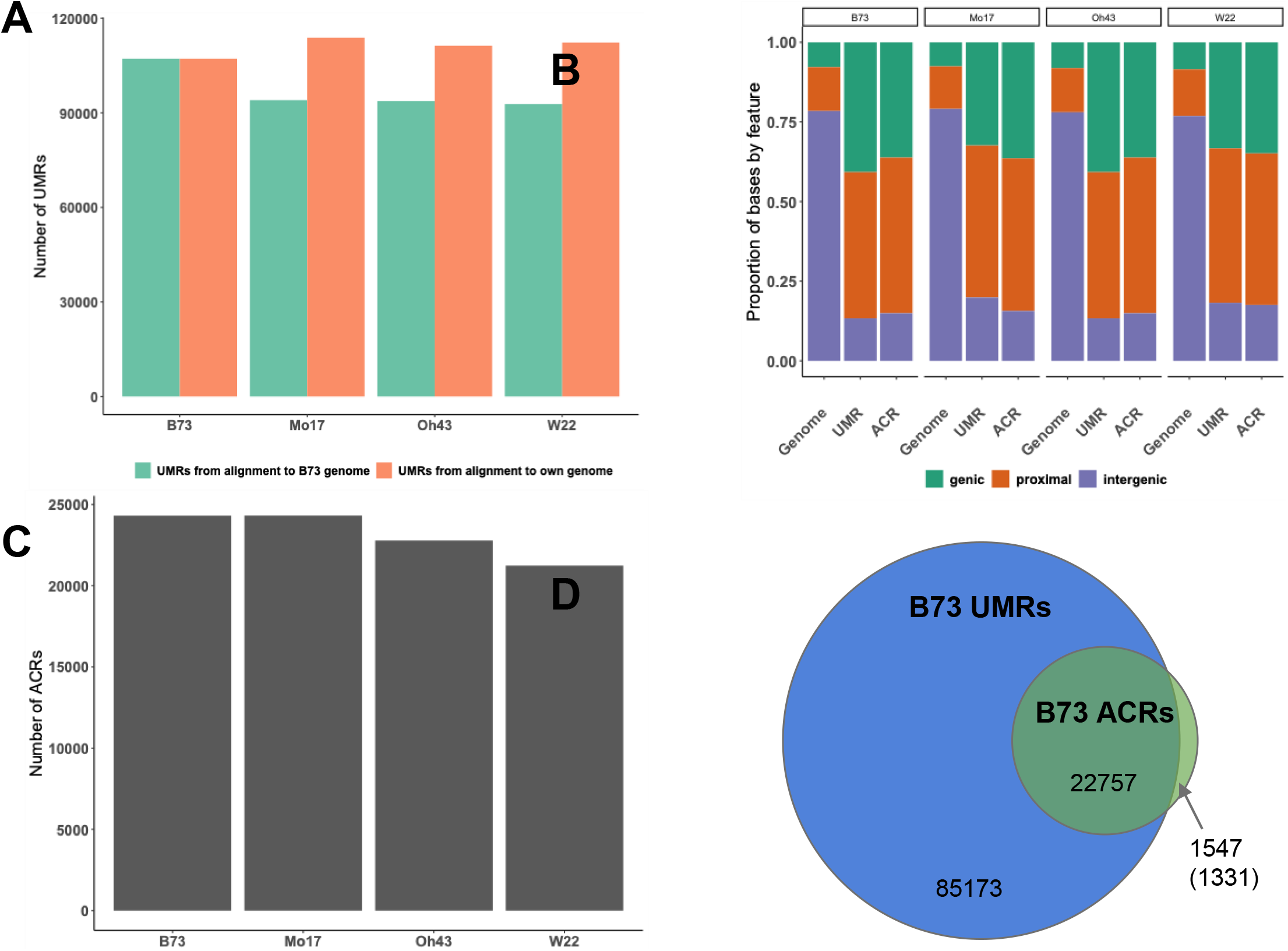
Identification of UMRs and ACRs in maize genotypes. A) The number of UMRs defined based on samples aligned to B73v4 (green) and their own genome assembly (orange). B) The location of UMRs and ACRs in the genome based on gene annotations was classified as overlapping genes (green), within 2kb of a gene (orange) and >2kb from a gene (purple). C) The number of ACRs defined based on the merged replicates for each genotype aligned to their respective genome assemblies. D) Overlap between the B73 UMRs and ACRs defined based on alignments to the B73v4 genome. Number in parentheses indicates ACRs that are defined as methylated as opposed to missing data.

Prior studies have found that unmethylated portions of the maize genome often contain cis-regulatory regions (Oka *et al.* 2017; Ricci *et al.* 2019; Crisp *et al.* 2020). To determine the concordance between UMRs and ACRs, we implemented ATAC-seq in the same four genotypes. ACRs were identified in each individual sample as well as from merged biological replicates (Table S2). We focused on analysis of the ACRs identified from the merged replicates, since the data from the two biological replicates was highly correlated and the ACRs identified within individual samples were frequently found in the merged sample (Table S3, Figure S2). There are 21,232-24,309 ACRs present in each of the four genotypes (Table 1, Figure 1C). Relative to UMRs, the ACRs are more enriched in gene-proximal regions of the genome and depleted within intergenic regions, but >24% of the ACRs are found >2kb from the nearest gene (Figure 1B). The vast majority of ACRs are found within UMRs in each of the four genotypes (Figure 1D, S3). While the vast majority of ACRs occur within UMRs, there are many UMRs without accessible chromatin (Figure 1D). This allows the classification of UMRs as accessible UMRs (aUMRs) or inaccessible UMRs (iUMRs) based on whether they overlap an ACR. The presence of an aUMR, which includes the presence of an accessible chromatin region, is much more common within or near genes that are highly expressed, but is quite rare for lowly expressed genes (Figure S3D). In contrast, iUMRs are present near genes with low and high expression levels, but are depleted near silent genes (Figure S3E). While the aUMRs represent an overlap between an unmethylated region and chromatin accessibility, the boundaries of these regions are often not the same. The majority (97.3%) of cases represent a larger unmethylated region in which the ACR only covers a portion of the UMR and the ACR is often found in the center of the UMR (examples in Figure S4). This suggests that the transition from accessible to inaccessible chromatin and from unmethylated DNA to methylated DNA does not occur at the same region.

### Classification of shared and non-shared genomic regions

Previous studies have assessed natural variation in DNA methylation based on alignment to a single reference genome (Regulski *et al.* 2013; Li *et al.* 2015b). However, when WGBS data from non-B73 genotypes are aligned to the B73 genome, the proportion of regions with missing data increases substantially (Figure S1B) and the methylation levels for genomic regions missing in B73 are not assessed. The availability of multiple reference genomes provides the opportunity to assess DNA methylation levels in the pan-genome that includes both shared (syntenic) regions of the genome with or without allelic variation, as well as non-shared regions that are present in one line and missing in another. The alignment of WGBS or ATAC-seq data to their respective genome provides the advantage of more complete characterization of DNA methylation and/or chromatin accessibility, but introduces complications for the direct comparison of specific regions among genomes.

To address this complication in comparing regions across genomes, chromosomal alignments were performed between the B73 genome and the other reference genomes to identify the shared and non-shared genomic segments between any two genotypes (see Methods) (Figure 2A). The approach that was implemented employed relatively stringent criteria for identification of shared regions. The regions classified as non-shared include both structural variants and highly polymorphic regions as well as highly repetitive regions that could not be uniquely mapped. Approximately 55% of the non-B73 genome sequences could be classified as syntenic and mappable relative to B73, with the remaining 45% not aligning to the B73 genome due to non-syntenic sequence or unmappable regions (Figure 2B). As a quality control measure, we assessed the proportion of space classified as shared or non-shared within identity-by-state (IBS) regions between genomes. The majority (94%) of IBS regions are classified as shared between any two genomes (Table S4) and the regions that are not classified as shared within IBS regions are highly enriched for repetitive sequences.

**Figure 2:**
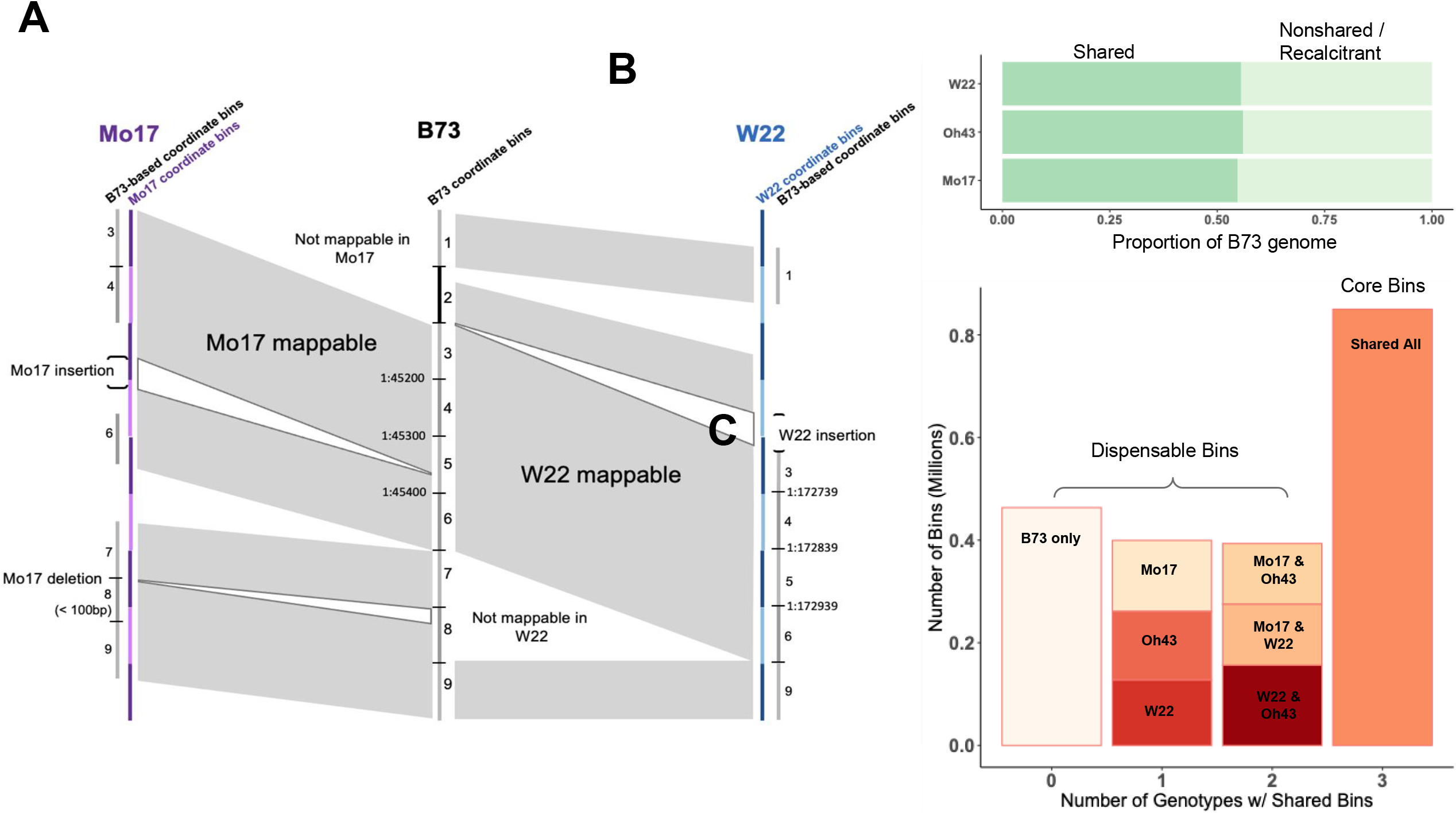
Defining shared and nonshared regions between genome assemblies. A) Schematic representation of B73-based 100bp bins defined as shared or nonshared in Mo17 and W22 (gray shaded regions) based on chromosomal alignments. The 100bp bins in W22 or Mo17 could be defined by 100bp increments within that genome sequence or based on coordinate matches to the B73 genome and these are shown as the W22 (blue) or Mo17 (purple) coordinate bins or the B73-based coordinates (grey). The black hash or the light to dark color change indicates the 100bp bin boundaries. B) The proportion of the B73 genome that is defined as shared or non-shared with Mo17, W22, and Oh43 based on chromosome-level sequence alignments. C) The number of B73 100bp bins that are unique to B73 (0 shared genotypes), shared with one other genotype assessed (1), shared with two other genotypes assessed (2) or shared across all 4 genotypes including B73, Mo17, Oh43, and W22 (3). Genotype labels correspond to the genotypes which share 100bp bins with B73.

Our analysis of DNA methylation or chromatin accessibility is often focused on 100bp bins. To directly compare the same coordinate space between genomes, we identified the 100bp bins from the B73 genome that were shared across genotypes (Figure 2A, S4). In the comparisons of B73 to the other three genomes, we find 41-48% of the B73 bins are non-shared, 37-42% of bins have an exact match in shared regions, 12-14% mapped with >= 1 SNP, and an additional 4% mapped with >= 1 small (<20bp) indel between the two genotypes. Across all comparisons, there are over ~800,000 100bp bins that are shared in all four genotypes (Figure 2C). There are ~500,000 bins that are found only in B73 and another ~800,000 that are present in B73 and only one or two of the other two genotypes (Figure 2C). The regions that are shared between genotypes have fewer bins with missing data such that only 6.7% of the bins shared in all three genotypes lack DNA methylation data compared to 28.4% of the bins that are only present in B73. This likely reflects the fact that much of the non-shared sequence between genomes is highly repetitive and recalcitrant to unique mapping. The identification of these shared bins allowed us to calculate the methylation levels or ATAC-seq read depth for the specific coordinates in a second genome that correspond to the B73 bins to allow direct comparisons of chromatin properties between genomes using epigenomic data aligned to its own reference genome.

### UMRs and ACRs are depleted in non-shared portions of the genome

We initially focused on the chromatin properties of the non-shared portions of the genome to assess the frequency of UMRs or ACRs within the dispensable portion of the genome compared to the shared portions. While over 10% of the shared genomic regions are annotated as genic less than 4.8% of the non-shared regions are annotated as genic reflecting a depletion of genes and enrichment of intergenic and TE sequence. The analysis of the *bronze1 (bz1)* locus on chromosome 9 illustrates these trends of shared space in genic regions and large non-shared blocks between genes, as previously described (Fu and Dooner 2002; Wang and Dooner 2006) (Figure 3A). In the *bz1* region, very few UMRs or ACRs are found within the non-shared regions (Figure 3B). We proceeded to perform a genome-wide assessment of the proportion of UMRs within shared and non-shared regions of the genome. While UMRs account for 6% of the entire B73 genome, only ~2% of the non-shared genomic regions are classified as UMRs compared to ~12% of the shared genomic regions (Figure 3C). A similar analysis of the genome-wide distribution of ACRs reveals that accessible chromatin is even more enriched within genomic regions that are shared among all four genotypes (Figure 3C). ACRs account for 1.2% of the shared genomic space but only 0.1% of the non-shared genomic regions (Figure 3C). Both ACRs and UMRs tend to be enriched near genes and the non-shared genomic regions are relatively gene-poor. However, this depletion of genes is not the only explanation for the paucity of ACRs and UMRs in the non-shared genomic regions. Over 80% of the genes in the shared space contain a UMR, while only 17% of the genes located in non-shared regions contain UMRs. Prior studies have found that non-shared genes are less likely to be expressed (Hirsch *et al.* 2016; Sun *et al.* 2018; Anderson *et al.* 2019; Haberer *et al.* 2020) and the depletion of UMRs within or near these genes further suggests that many of these “genes” lack the chromatin properties associated with expression. These analyses suggest that pan-genome assessment of UMRs and ACRs will provide a more complete identification of UMRs/ACRs but that there are a limited number of novel UMRs or ACRs in non-shared space in maize. The subsequent analysis will focus on the UMRs and ACRs that are present within shared regions of any two maize genomes.

**Figure 3:**
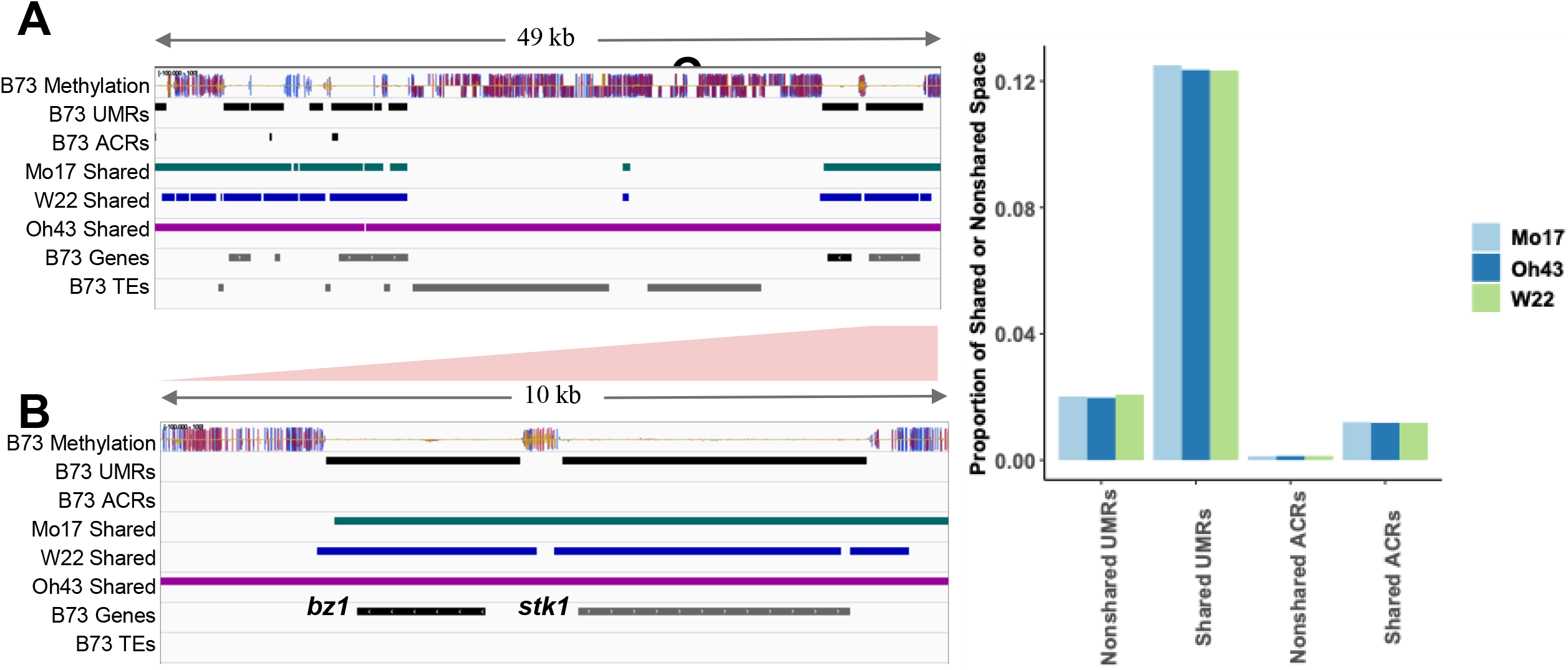
Presence of ACRs and UMRs within shared and non-shared genomic regions. A) An IGV (Robinson et al., 2011) representation of a 49kb segment on chromosome 9 of the B73 genome assembly. Tracks show B73 methylation levels in all contexts (CG-blue, CHG-red, CHH-yellow), B73 UMRs and ACRs, Mo17 shared sequence (green), W22 shared sequence (blue), Oh43 shared sequence (purple), and B73 gene and TE annotations (grey). B) A small region of the bz1 locus was expanded to see the detail. C) The B73 genome was compared to Mo17, Oh43 or W22 to define regions that are shared or non-shared in each contrast. The proportion of the shared or non-shared space that is classified as UMR or ACR was determined for each of the pairwise contrasts.

### Comparisons of UMRs and ACRs in the shared space of maize genomes

We proceeded to focus on the UMRs and ACRs that are present within shared regions between maize genomes. The analyses were primarily focused on UMRs as these encompass the vast majority of ACRs (Figure 1D) and we could monitor stability for the UMRs with an ACR (aUMRs) compared to the UMRs without an ACR (iUMRs). The B73 UMRs were compared to each of the other genomes and classified based on whether they are present in shared/non-shared regions and then whether the region has DNA methylation data available for both genotypes. For the ~90% of B73 UMRs that have defined methylation states and are present in a shared region we could classify whether there is an overlapping UMR in the other genotype or whether the UMR is polymorphic such that it is classified as methylated in the other genotype (Figure 4A). Most UMRs that are present in shared space overlap a UMR in the other genotype while a small set are polymorphic (Figure 4A). The overlapping UMRs can be classified as identical if the boundaries of the UMR are the same in both genotypes (example in Figure 4B). Alternatively, an overlapping UMR could represent a partial overlap such that one genotype has a larger region than the other or both edges are shifted (examples in Figure 4B). The UMRs with partial overlap account for the majority of the overlapping UMRs between two genotypes (Figure 4A).

**Figure 4:**
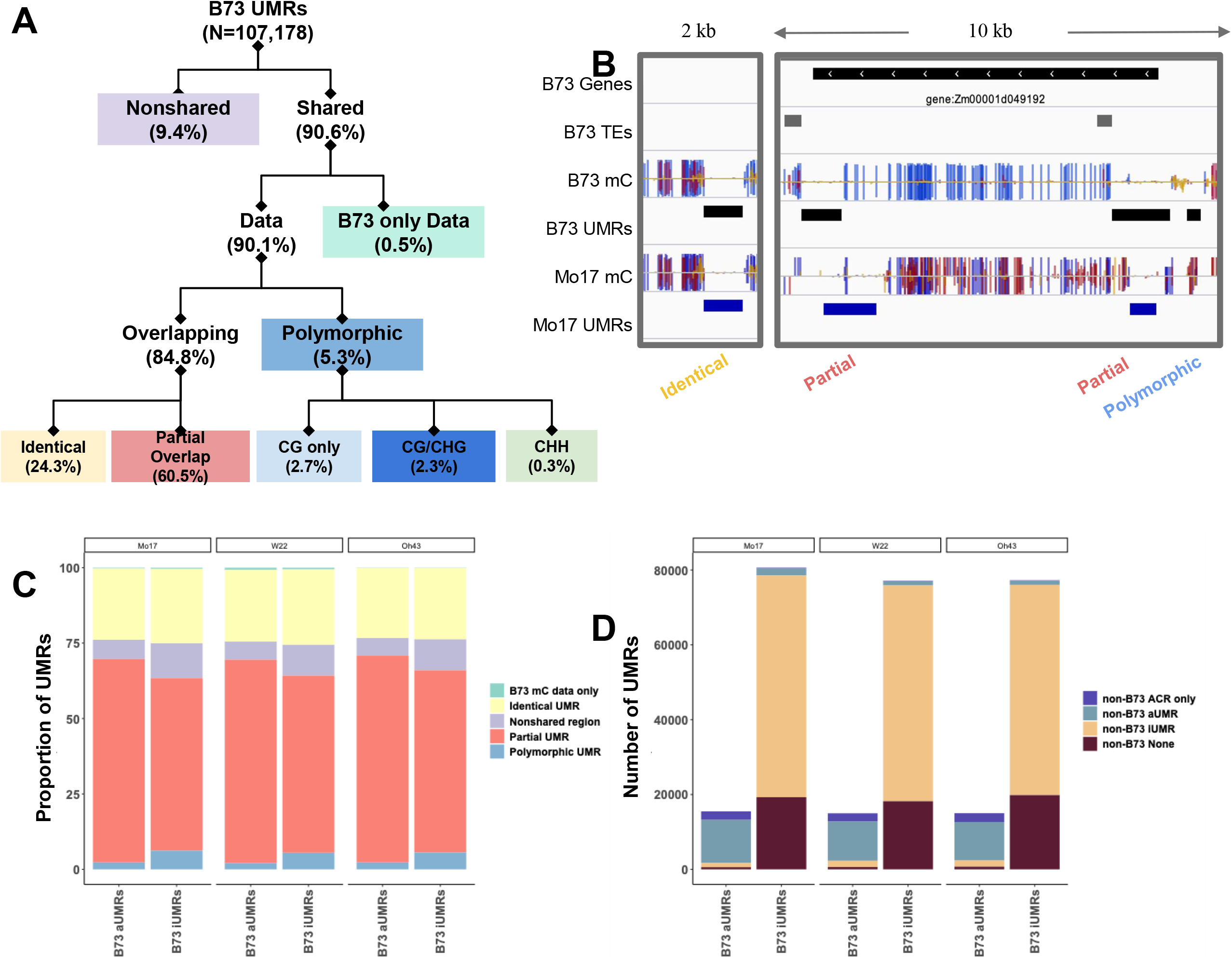
Stability of UMRs in shared sequence. A) A flowchart on how B73 UMRs are classified is shown. The numbers in parenthesis indicate the average number of regions classified in that group based on comparisons to the other genotypes. The proportion of B73 UMRs that are shared or non-shared (purple) based on sequence with the respective genome assembly. Shared regions are further classified as B73-only (green) for UMRs that lack data in the other genotype, identical (yellow) for UMRs that maintain an unmethylated state in the same region, partially overlapping (pink) for UMRs that maintain an unmethylated state but have different UMR boundaries across genotypes or polymorphic (blue) for UMRs that change to a methylated state in the other genome. The colors in A are identical to those in C. B) A genome browser view of the several regions in the B73 genome to illustrate examples of identical, partially overlapping and polymorphic UMRs. A track of DNA methylation in all contexts (CG-blue, CHG-red, CHH-yellow) is shown for B73 and Mo17 (both aligned to B73v4) with UMRs defined below in black (B73) and blue (Mo17). B73 UMRs are defined as identical (yellow), partial overlap (pink), or polymorphic (blue). C) The proportion of B73 UMRs that are classified in each group defined in A are shown for both aUMRs and iUMRs based on comparison to each of the other three genotypes. D) The number of B73 aUMRs or iUMRs that are classified as ACR only (not unmethylated) in the other genotype (purple), aUMR in the other genotype (blue), iUMR in the other genotype (yellow), or methylated and inaccessible in the other genotype (burgundy) are shown for comparisons to each of the other genotypes

B73 UMRs can be subdivided into aUMRs and iUMRs based on the presence, or absence, of an ACR within the unmethylated region. We compared the distribution of classifications for the aUMRs and iUMRs for the presences of identical, partially overlapping, or polymorphic UMRs in the other genotypes (Figure 4C). The B73 aUMRs have fewer examples of polymorphic UMRs as well as fewer examples within non-shared genomic regions. However, this is largely due to a larger proportion of overlapping UMRs that are partially overlapping rather than more examples of identical UMRs (Figure 4C). These analyses suggest that while any two genomes often have UMRs in similar regions the exact coordinates of the UMRs are often distinct.

The B73 aUMRs and iUMRs were also assessed for the potential changes to either methylation or accessibility between genotypes (Figure 4D). The majority (~71.2%) of the B73 aUMRs were maintained as aUMRs in the other genotypes. However, there are also a subset of the B73 aUMRs that lose either the unmethylated state (~14.8%) or chromatin accessibility (~11.1%) in the other genotype. The remaining 2.9% are not classified as either ACR or UMR for the same region in the other genome. The B73 iUMRs often (~73.7%) are unmethylated and inaccessible in the other genotypes (Figure 4D). There are also many (~24.5%) examples of B73 iUMRs that are methylated in the other genotype. The proportion of shifts from unmethylated to methylated states are much higher for the iUMRs than the aUMRs. Very few (~1.6%) of the B73 iUMRs exhibit accessibility in the other genotypes (Figure 4D).

### Unique properties of regions with methylation changes in various methylation contexts

While the polymorphic B73 UMRs that are methylated in another genotype only account for a small set of all UMRs these may represent important functional differences between genotypes. The polymorphic UMRs can be subdivided based on the prominent class of methylation in the other genotype (Figure 4A, 5A). Each of these classes of methylation gains likely reflect distinct mechanisms and chromatin types. The types of methylation observed in these regions do not reflect the genome-wide proportions of methylation types (Figure S1). The proportions that are classified as CG only or CHH are higher than observed genome wide (Figure S1, 5B). While CG/CHG regions are depleted, although there are still many examples of CG/CHG at these regions of variable methylation (Figure 5B).

**Figure 5:**
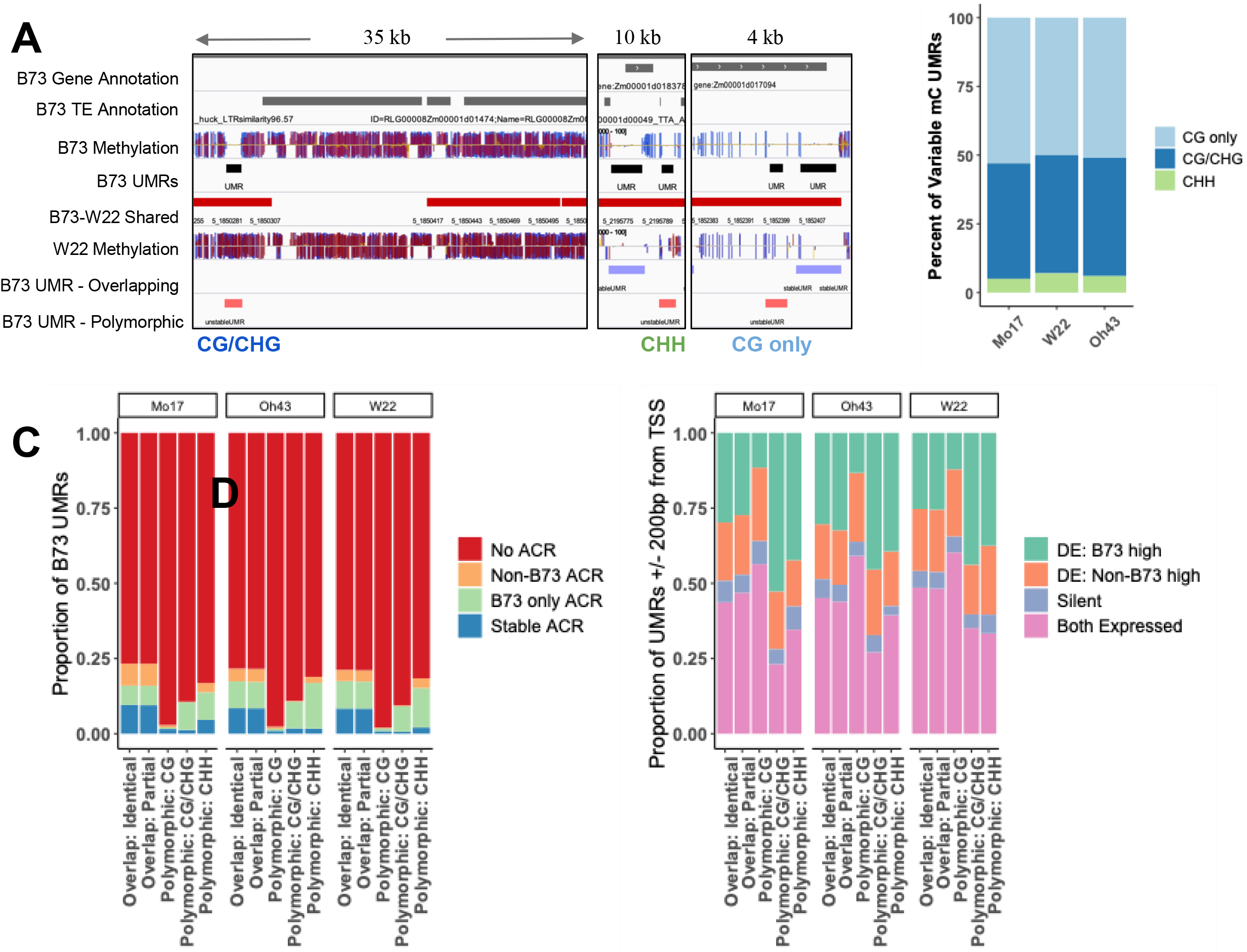
Characteristics of polymorphic UMRs. All B73 UMRs classified as polymorphic (shown in Figure 4A) were assessed based on the type of methylation present in the methylated genotype. The classification is based on which type of methylation state is most common among the 100bp bins of the UMR. A) A genome browser view of a region on chromosome 5 of the B73 genome. A track of B73 methylation in all contexts (CG-blue, CHG-red, CHH-yellow) is shown with UMRs defined below in black. Regions with shared sequence with W22 are shown in red and the W22 methylation track (aligned to the B73v4 assembly) with corresponding UMR classification as overlapping (purple) or polymorphic (red). Three separate snapshots are shown with the type of methylation found in W22 for the variable UMR noted below (CG only, CG/CHG, or CHH). B) The percent of all B73 UMRs classified as polymorphic that change to CG only (light blue), CG/CHG (dark blue), or CHH (green) methylation in the other genotype was calculated. C) UMRs were defined as containing an ACR in both genotypes (Stable ACR: blue), in one genotype (B73 only ACR: green, Non-B73 ACR: orange), or lacking an ACR in both genotypes (No ACR: red). The proportion of each category of B73 UMR (overlapping and polymorphic) that is defined by ACR presence or absence is shown for each genotype. D) The proportion of UMRs that are found within 200bp of an annotated gene TSS that are defined as differentially expressed (DE), expressed in both genotypes or not expressed is shown for each genotype. Genes were classified as differentially expressed (log2 fold change > 2 and p-value < 0.05) with the higher expression level observed in B73 (green) or the non-B73 genotype (orange) or as non-differentially expressed (FPKM > 1, pink) or not expressed (silent: purple).

The presence or absence of ACRs in both genotypes was assessed for the polymorphic UMRs relative to overlapping UMRs (Figure 5C). While both identical UMRs and partially overlapping UMRs show virtually identical proportions with shared ACRs or polymorphic ACRs, the polymorphic UMRs have very few stable ACRs (Figure 5C). This is expected as there are very few examples of accessible regions within methylated DNA. As expected, there are very few examples of ACRs only in the methylated genotype. The proportion of the polymorphic UMRs that are classified as having an ACR only in B73 but not the methylated genotype is quite variable. Polymorphic UMRs with CHH methylation in the other genotype are more likely to have an ACR in B73 than polymorphic UMRs with CG only methylation in the other genotype (Figure 5C). This could reflect the fact that CHH methylation is often found in regions immediately upstream or downstream of genes in the maize genome (Gent *et al.* 2013; Li *et al.* 2015a) and that these regions often have ACRs. In contrast the CG-only methylation often occurs within gene bodies, where ACRs are less common than at the edges of genes or promoter regions.

We proceeded to assess variable gene expression of genes near overlapping or polymorphic UMRs using RNAseq data from the same tissue used to monitor accessibility and DNA methylation. Genes with an overlapping or polymorphic UMR within 200bp (upstream or downstream) of the transcription start site (TSS) were identified and classified as being differentially expressed (DE), expressed in both genotypes but not DE, or silent (FPKM < 1 in both genotypes). Genes that have identical or partially overlapping UMRs near the TSS exhibit nearly identical proportions of genes in these categories and have similar proportions of genes that are higher expressed in B73 or the other genotype (Figure 5D). Polymorphic UMRs that gain CG-only methylation in the other genotype have fewer examples of genes with higher expression in B73. This suggests that the presence of CG only methylation is rarely associated with reduced gene expression. In contrast, genes with gains of CG/CHG or CHH methylation near the TSS are enriched for genes that are higher expressed in B73. While there is an enrichment for DE expression with the unmethylated genotype being higher expression for polymorphic UMRs with CG/CHG or CHH methylation gains, it is worth noting that there are also many examples of genes near these types of polymorphic UMRs that are not differentially expressed. This suggests that the gain of CG/CHG methylation or CHH methylation in regions surrounding the TSS can be associated with altered expression in some cases, but that other genes can tolerate variable methylation without a significant change in expression.

### Partially overlapping UMRs contribute substantially to differentially methylated regions

The analysis of natural variation for DNA methylation is often focused on identification of differentially methylated regions (DMRs) between genotypes. In this study, we elected to focus on the conservation / variation of unmethylated regions as these regions have evidence for functional relevance in crop genomes. However, the observation that many of these regions only have partial overlap suggests that many DMRs might be the result of a shift in the boundary between methylated and unmethylated DNA rather than a complete regional gain/loss of methylation (Figure 6A). The 100bp bins were used to identify DMRs between the genotypes. There are 116,000-158,000 100bp bins that are classified as differentially methylated with hypomethylation in B73 relative to the other genotype. We assessed how many of these DMRs are due to completely polymorphic UMRs compared to partial UMRs with different boundaries between methylated and unmethylated DNA (Figure 6B). The polymorphic UMRs account for 2.5-3.3% of all differentially methylated bins depending on which genotypes are being compared. A larger proportion (51.5-53.5%) of the differentially methylated bins are due to partially overlapping UMRs. The remaining differentially methylated bins occur in regions too small to be classified as UMRs (unmethylated regions <300bp) or represent single bin differences in larger UMRs. This analysis suggests that many of the DMRs are due to shifting boundaries between methylated and unmethylated DNA rather than a complete gain or loss of methylation in a region.

**Figure 6:**
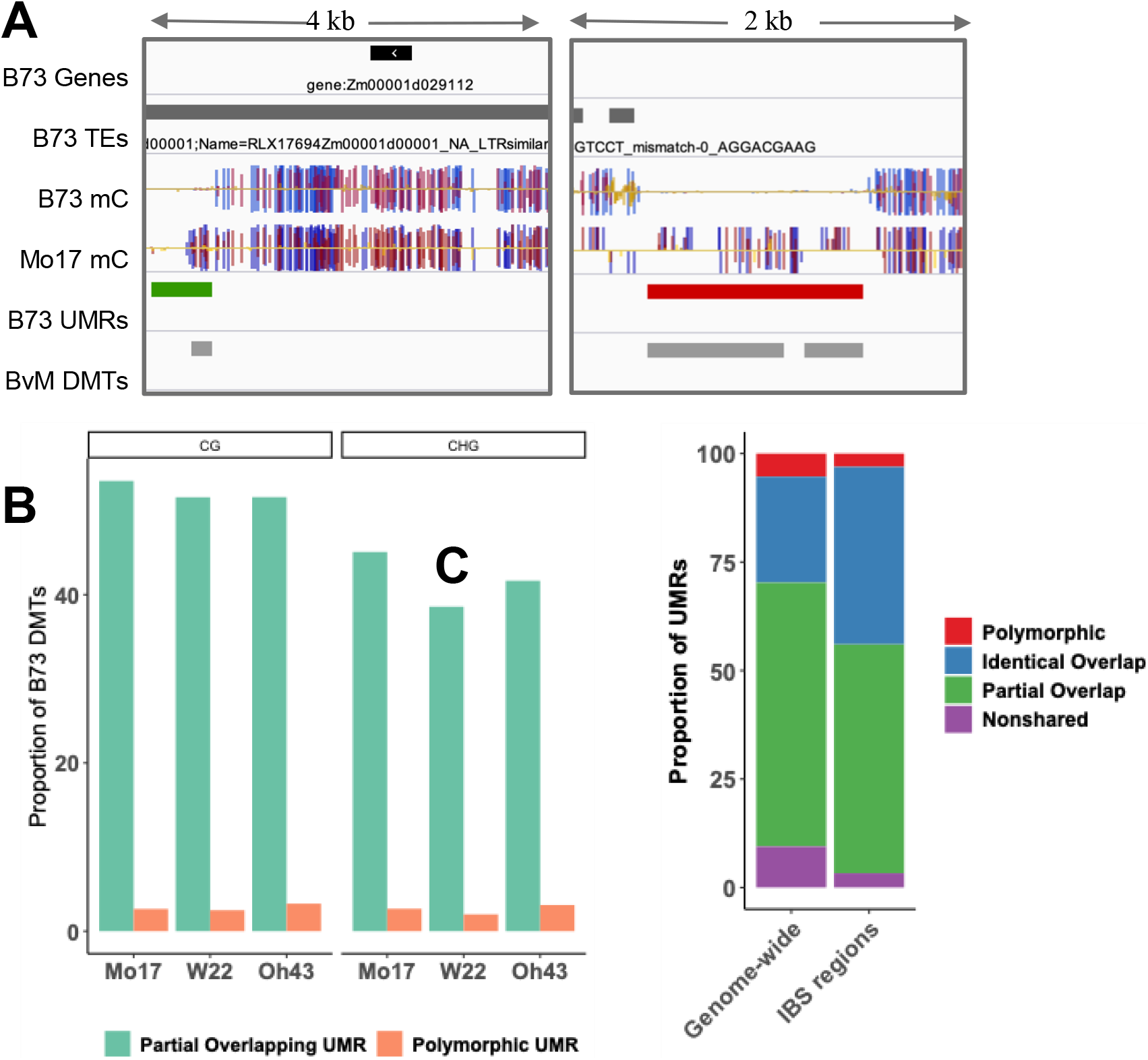
Many differentially methylated tiles (DMTs) are due to partially overlapping UMRs. A) IGV (Robinson et al., 2011) view of DMTs. Tracks show B73 gene and TE annotations, B73 and Mo17 single cytosine methylation in all contexts (CG: blue, CHG: red, CHH: yellow), B73 UMRs and classification relative to Mo17 (identical: blue, partial: green, polymorphic: red), and DMTs defined by a low level of B73 CG methylation and high level of Mo17 CG methylation. B) The proportion of B73 DMTs that are associated with partially overlapping UMRs (green) or polymorphic UMRs (orange) is shown. C) The proportion of B73 UMRs, genome-wide (control) or in IBS regions, that are shared or non-shared (purple) based on sequence with the respective genome assembly. Shared regions are further classified as missing data (orange) for UMRs that lack data in the other genome, identical (blue) for UMRs that maintain an unmethylated state in the same region, partially overlapping (green) for UMRs that maintain an unmethylated state but have different UMR boundaries across genotypes or polymorphic (red) for UMRs that change to a methylated state in the other genome.

These observations suggest that the specific boundary between methylated and unmethylated DNA can be variable between genotypes. This could be due to sequence changes at or near the edges of these regions or could arise due to stochastic variation with no sequence change. To address this question we assessed the proportion of identical or partially overlapping UMRs within large (>1Mb) blocks of sequence that is IBS. In total there was 112.7Mb of IBS sequence blocks that could be assessed and these are large blocks of sequence that are essentially devoid of SNPs or structural variants. Within these regions we find a depletion for polymorphic UMRs. While 5.3% of all UMRs are classified as polymorphic we find only 2.8% of UMRs that are classified as polymorphic in these regions suggesting that fully polymorphic UMRs are depleted in the absence of sequence variation (Figure 6C). The IBS regions have a higher proportion of UMRs with identical boundaries in the two genotypes. However, there are still a large number of UMRs with shifted boundaries (49.5%) suggesting that the boundaries between methylated and unmethylated DNA can shift even without nearby sequence variation.

## Discussion

*Zea mays,* unlike many other model organisms, has a large genome containing 80% repetitive sequence and high levels of DNA methylation interspersed with functional genic and regulatory regions (Schnable *et al.* 2009; Jiao *et al.* 2017). Examination of genome structure across inbred lines have identified extensive polymorphism in both genic and repeat regions of the maize genome (Chia *et al.* 2012; Hirsch *et al.* 2014; Springer *et al.* 2016; Darracq *et al.* 2018; Anderson *et al.* 2019; Hufford *et al.* 2021). Prior analyses of natural variation of chromatin in maize have been based on epigenome profiling data aligned to a single reference genome (Li *et al.* 2015b; Xu *et al.* 2020). While a single reference genome provides insight into variation in conserved genomic regions, it does not contain the full set of sequences present in the lines being compared, resulting in biases in the ability to compare chromatin properties. The availability of multiple *de novo* genome assemblies allows for a more complete discovery of regions with specific chromatin properties, such as UMRs or ACRs. In this study, we profiled genome-wide DNA methylation, based on alignments of data to the corresponding genome assembly, to identify the ~6% of each genome that exhibits an unmethylated state and the ~1% that is accessible chromatin. A pan-genomic analysis of UMRs and ACRs reveals the frequency of these features within both shared and nonshared genomic regions. Within the shared sequence regions it is possible to assess the stability of the unmethylated and accessible chromatin portions of the genome.

### Pan-genome analyses reveal enrichment of unmethylated regions within shared sequence

Whole genome alignments between B73 and Mo17, W22, and Oh43 allowed for the identification of both shared and nonshared sequences. In a comparison of any two genomes, the sequence unique to each genome is primarily composed of highly repetitive sequences with extensive DNA methylation and is found to be depleted for genes (Chia *et al.* 2012; Springer *et al.* 2016; Hirsch *et al.* 2016; Darracq *et al.* 2018; Anderson *et al.* 2019; Hufford *et al.* 2021). The proportion of the nonshared genome that is classified as UMR or ACR is 6-12 fold lower than the proportion of the shared genome classified as UMR or ACR. This reduction in UMRs and ACRs is not simply due to the reduced gene content in nonshared space. Most (80%) of genes in the shared space are associated with a UMR, while only 17% of genes in non-shared space have a UMR. This is not unexpected as prior studies of presence-absence variation (PAV) genes have found that most of these genes that vary between genotypes are not expressed even when they are present (Swanson-Wagner *et al.* 2010). A recent analysis of 26 maize genomes that used a slightly different approach to classify unmethylated and CG-only regions reported similar findings (Hufford *et al.* 2021). More UMRs are identified by aligning chromatin data to the proper genome but the proportion of UMRs or ACRs in this nonshared space is much lower than in the shared regions. These analyses suggest that pan-genomic analyses can identify novel UMRs or ACRs but that these are relatively rare in the sequences that exhibit large scale structural variation. However, it is worth noting that the UMRs or ACRs that are present near the genes in the non-shared space can be an effective tool for identifying genes with potential expression (Sartor *et al.* 2019; Crisp *et al.* 2020). Given that many of the genes within these regions are likely pseudogenes generated by transposition of genes or gene fragments that can be difficult to annotate just based on sequence, the use of chromatin data can help to identify genes with potential function in these regions.

### Characterization of relative dynamics of accessibility and methylation

We were interested in studying the relative dynamics of both DNA methylation and chromatin accessibility among genotypes. Prior studies have found that the majority of accessible regions have little or no methylation (Ricci *et al.* 2019) but that there are also many unmethylated regions that lack accessibility (Crisp *et al.* 2020). The analysis of UMRs that are present within shared sequence regions can be used to understand how often there is variation in only accessibility as opposed to coordinate changes in both DNA methylation and accessibility. The accessible UMRS (aUMRs) tend to be relatively stable in other genotypes with both accessibility and lack of DNA methylation for an overlapping region in other haplotypes. This is consistent with the concept that these regions may be important for proper regulation of gene expression and therefore changes in these chromatin properties could be associated with functional differences. The inaccessible UMRs (iUMRs) were often inaccessible and unmethylated in both genotypes but there were a large number of these that exhibit polymorphic DNA methylation status such that they exhibit high levels of DNA methylation in the other genotypes. Only a small proportion of these UMRs exhibit a consistently unmethylated state in both genotypes with accessible chromatin in only one of the two genotypes. These likely include some examples of false negatives due to relatively stringent criteria for calling an ACR. In these cases, an ACR may be present in both genotypes but only identified as significant for one genotype. However, these cases of variable chromatin accessibility also include examples with clear support for chromatin accessibility in one genotype but no evidence for chromatin accessibility in the other genotype. These are interesting as they potentially reflect differences in transcription factor occupancy for regions that are stably unmethylated in both genotypes. It is possible that these may reflect differences in tissue-specific expression patterns of some maize genes. In leaf tissue there may be differential chromatin accessibility, but it is possible that the genotype without chromatin accessibility in leaf tissue still becomes accessible in some other tissue that exhibits expression. Alternatively, minor sequence changes at transcription factor binding sites may result in loss of chromatin accessibility even though the region is unmethylated in both genotypes.

### Stability and instability of UMRs between genotypes

A subset of the shared sequence UMRs do not maintain their unmethylated state across genotypes and instead have high levels of methylation in at least one of the other three genotypes. The presence of methylation variation in the shared sequence regions allowed for characterization of attributes associated with chromatin state instability. Prior studies have suggested that structural variants, especially transposable element polymorphisms, can be associated with changes in DNA methylation for nearby sequences (Eichten *et al.* 2012; Schmitz *et al.* 2013). When analyzing DNA methylation based on a single reference genome it can be difficult to incorporate information about structural variants and to map reads near the junctions of these variants. Using alignments to each reference genome and then comparing coordinates of syntenic 100bp tiles allowed us to monitor changes in DNA methylation between genotypes, even in regions near structural variants. The polymorphic UMRs that represent a full shift of an unmethylated region in one genotype to methylation in the other genotype are depleted in regions devoid of structural variants. Within large blocks of IBS 2.8% of the UMRs are polymorphic. In contrast, over 5.3% of all UMRs are classified as polymorphic. This indicates that changes in methylation state can occur in the absence of nearby structural variants but that the rate is substantially higher in regions with sequence variation.

In this study, we focused on the conservation and variation for UMRs or ACRs between genotypes. These are relatively large (at least 300bp based on the criteria used for discovery) regions that lack DNA methylation. We focused on these regions due to prior evidence for functional enrichment of these regions (Oka *et al.*; Ricci *et al.* 2019). We note that most of the UMRs in one genotype have an overlapping UMR in another genotype. This suggested stability of these chromatin patterns among genotypes. However, closer inspection revealed that the majority of these overlapping UMRs have different boundaries in the two genotypes. These include examples in which one UMR is entirely within the other as well as examples that have partial overhangs in both genotypes. The partially overlapping UMRs seem to have very similar genomic distributions and overlap with ACRs or altered gene expression in similar proportion to those for identical conserved UMRs. This suggests that these shifts in the boundary between methylated and unmethylated DNA do not have functional impact in most cases. This may suggest that the presence of a UMR is more defined by sequences in the middle of the unmethylated region rather than particular sequences at the edges that define the extent of methylation.

The observation of many partially overlapping UMRs suggested that these shifts in the boundary between methylated and unmethylated DNA could account for many examples of differential methylation between genotypes. Conceptually, it is tempting to think that most differentially methylated regions result from a local gain or loss of a patch of DNA methylation. However, our analyses suggest that many of the differentially methylated 100bp tiles actually arise due to changes in the boundaries between UMRs in different genotypes. Further studies will be necessary to determine if these differences in methylation boundaries represent a continuum such that each genotype has a slightly different boundary or if there are preferred epi-haplotypes.

## Methods

### Reference Genomes

Whole genome assemblies for four maize inbred lines, B73 (Jiao *et al.* 2016), W22 (Springer *et al.* 2018), Mo17 (Sun *et al.* 2018), and Oh43 (Hufford *et al.* 2021) were used for genome-wide analyses. All analyses were performed on assemblies of chromosomes 1-10 while all unplaced scaffolds were disregarded due to the inability to compare these regions across genotypes. Filtered gene and structural TE annotations (Stitzer *et al.*; Anderson *et al.* 2019) were used.

### Sample Collection

Maize B73, W22, Mo17 and Oh43 plants were grown under 16 h/8 h 30°C /20°C day/night for 13 days in the growth chamber of University of Minnesota. DNA was extracted from leaves of two-week old V2 plants using the DNeasy Plant Mini kit (Qiagen). Four or five biological replicates consisting of a pool of tissue from 4 plants were collected for each genotype. Two of these biological replicates were sampled for profiling of DNA methylation and chromatin accessibility while all biological replicates were used for RNAseq.

### WGBS protocol

Two technical replicates of each genotype (B73, Mo17, W22, and Oh43) were generated. 1ug of DNA in 50ug of water was sheared using an Ultrasonicator to approximately 200-350bp fragments. 20ul of sheared DNA was then bisulfite converted using the EX DNA Methylation-Lightning Kit (Zymo Research) as per the manufacturer’s instructions and eluted in a final volume of 15ul. Then 7.5ul of the fragmented bisulfite-converted sample was used as input for library preparation using the ACCEL-NGS Methyl-Seq DNA Library Kit (SWIFT Biosciences). Library preparation was performed as per the manufacturer’s instructions. The indexing PCR was performed for 5 cycles. Libraries were then pooled and sequenced on a NovaSeq 6000 in high output mode 125bp paired end reads over a single lane at the University of Minnesota Genomics Center. WGBS data generated in this study is deposited at NCBI SRA and available under accession.

Trim_glore(Martin 2011) was used to trim adapter sequences and read quality was assessed with the default parameters in paired-end read mode plus a hard clip of 20bp on each read due to SWIFT protocol specifications. Reads that passed quality control were aligned to their corresponding genome assemblies. Alignments were conducted using BSMAP-2.90(Xi and Li 2009), allowing only unique hits with up to 5 mismatches and a quality threshold of 20 (-v 5 -q 20). Duplicate reads were detected and removed using picard-tools-1.102 (“Picard”) and SAMtools(Li *et al.* 2009). Conversion rate was determined using the reads mapped to the unmethylated chloroplast genome. The resulting alignment file, merged for all samples with the same tissue and genotype, was then used to determine methylation level for each cytosine using BSMAP tools.

### Methylation data summary

Methylation levels were summarized using the bsmap methratio.py script to group by context (CG, CHG, CHH). The number of cytosines in every 100bp bin of the genome was determined and the proportion of cytosines defined as methylated was calculated. Coverage was calculated as CT / # of sites for each context. Methylation domain was classified for each 100bp bin based on the protocol described in Crisp et al. (Crisp *et al.* 2020) with criteria defined as a minimum site count of 2 and coverage of 3. UMRs were defined by grouping adjacent unmethylated bins.

### ATAC-seq protocol and ACR classification

ATAC-seq libraries were generated as described in Lu et al (Lu *et al.* 2017). Two technical replicates of each genotype (B73, Mo17, W22, and Oh43) were generated from the same samples as those used for WGBS data generation. Raw reads per sample were preprocessed with Trim_gloare. Trimmed reads were aligned to the *Zea mays* B73v4 genome and the genome assembly specific to each sample using Bowtie v1.2.3 with the following parameters: “bowtie -X 1000 -m 1 -v 2 --best –strata”. Aligned reads were converted to bam files and sorted using SAMtools v1.9. Clonal duplicates were removed using Picard MarkDuplicates v2.23.3 (http://broadinstitute.github.io/picard/). Input data of maize B73 was retrieved from a previous publication and processed to obtain bam files with clonal duplicates removed. MACS2 was employed to call initial ACRs with Input data as control (-c) and sample data as treatment (-t) using the following parameter “-g 2.1e9 --keep-dup all --nomodel --extsize 147”. The post-processing followed the same procedure as a prior publication (Ricci *et al.* 2019) to produce high-confident ACRs. Specifically, 1) Initial ACRs were split into 50 bp windows with 25 bp steps; 2) the Tn5 integration frequency in each window was calculated and normalized to the average frequency in the total genome; 3) windows with the normalized frequency greater than 25 were merged together allowing 150 bp gaps; 4) only merged regions greater than 50 bp were retained; 5) the mitochondrial or chloroplast genome from NCBI Organelle Genome Resources were removed using blast again sequences within merged ACR regions. The sites within ACRs that had the highest Tn5 integration frequency were defined as summits.

### RNA-seq protocol

RNA-seq data were generated in 150bp paired-end mode using NovaSeq 6000. B73, W22 and Mo17 reads were retrieved from the NCBI SRA accession PRJNA657262 (Liang et al. 2021) and Oh43 reads were deposited into NCBI SRA accession PRJNA692023. All of the raw reads were preprocessed using Trim_galore and aligned against the B73 AGPv4 reference genome using HISAT2 v2.1.0 (Kim *et al.* 2015). Gene annotations and disjoined TE annotations were used as described above. Gene exon regions were subtracted from TE regions and then appended to original TE annotation to remove ambiguous mapping between genes and TEs. Reads per gene or TE was determined using HTSeq-count v0.11.2 (Anders *et al.* 2015) and raw count data was input into DESeq2 (Love *et al.* 2014) to identify differentially expressed genes or TE elements.

The mean value for each feature (gene or TE) was calculated from 4 or 5 replicates. Any feature with a mean value greater than 1 was considered “expressed”. UMRs were associated with genes and TEs based on location relative to the feature. B73 UMRs which overlapped the annotated sequence coordinates within the genome being assessed were classified as “genic” or “TE”. Those not overlapping a gene but within 2kb of the gene start or end were classified as “proximal”.

### Cross-genotype mapping

Genome sequence from Mo17, W22 and Oh43 was first aligned to the B73 reference (Jiao *et al.* 2017) using minimap2 (Li 2018). The resulting alignments were merged and cleaned (removing overlapping alignment blocks and alignment blocks containing assembly gaps) using in-house perl scripts. BLAT Chain/Net tools were then used to create a single coverage best alignment net between the query genome (one of Mo17, W22 and Oh43) and the target genome (B73). Finally, a genome-wide synteny chain file was built for each genotype (against HM101), enabling downstream analyses such as variant detection and 100-bp tile liftover. Alignment pipeline and scripts are available on Github (https://github.com/baudisgroup/segment-liftover). Sequence was extracted for all 100bp bins in the B73 genome and aligned to Mo17, W22, and Oh43. Each bin was determined to be unmappable or mappable. Mappable bins were assigned coordinates in the non-B73 genome. The number of single nucleotide polymorphisms and insertion/deletions for each bin was calculated. Across all genotypes, only 4% of bins were found to have >= 1 insertion/deletion and 13% contained >=1 single nucleotide polymorphism. Bins with no more than 4 insertion/deletions of 20bp in size were kept for analyses of shared space. Each 100bp bin in B73 was designated as unmapped or provided matching sequence coordinates in each of the 3 other genotypes (Mo17, W22, Oh43).

### Differentially methylated tiles (DMTs)

WGBS data aligned to the respective genome and summarized in the B73-based 100bp coordinate system was used. Tiles were subset to those with sequence mappability and coverage in both genotypes for each pairwise comparison. DMTs were defined by a difference of 40% with at least one genotype having <10% and >40% methylation for CG and CHG contexts. CHH DMTs were defined by one genotype with <5% and >25% methylation in the 100bp tile. DMTs in each context were determined for Mo17, W22, and Oh43 compared to B73.

### Classification of UMR variability

B73 UMRs that were mappable to sequence in another genotype were further defined by methylation state in the corresponding genome. All 100bp bins within a defined UMR were assessed for the matching sequence coordinates in Mo17, W22, and Oh43. For each UMR, the proportion of bins classified as methylated (including CG, CG/CHG, and CHH methylation domains) was calculated. UMRs with >50% of the bins being methylated were defined as “polymorphic UMRs” for the difference in methylation state from unmethylated in B73 to methylated in the non-B73 genotype. All other UMRs, showing an unmethylated state in both B73 and the non-B73 genotype assessed, were defined as “overlapping UMRs”.

B73 UMRs that are methylated in another genotype (polymorphic UMRs) were further classified by the type of methylation observed in the non-B73 genotype. The polymorphic UMRs were summarized by domain. The proportion of 100bp bins with a methylated domain, within the defined B73 UMR, for each methylation context was determined. Any UMR that had >50% of its methylated bins classified as a specific methylation context was declared to be variable in that context. Classification was determined first by CHH methylation, followed by CG/CHG methylation and lastly CG only methylation. Variable methylation type was defined individually for each genome based on the sequence coordinates of the B73 UMR.

B73 UMRs that are unmethylated in another genotype (overlapping UMRs) were further classified by the coordinates of the defined UMR between B73 and the non-B73 genotype. The UMRs, defined by alignment of WGBS data to the B73 reference genome, were determined and their coordinates were assessed. Pairwise comparisons were done between B73 and non-B73 genotypes. B73 UMRs that had identical 100bp bin boundaries for the defined UMR were classified as identical UMRs. B73 UMRs that had variable boundaries were classified as partial UMRs (the coordinates of the smaller UMR were maintained within the larger UMR coordinates or the coordinates are shifted and have uniquely defined unmethylated bins in each genotype).

### Classification of ACR variability

Every B73 UMR was classified based on the accessibility of that shared sequence region within B73, Mo17, W22, and Oh43. All UMRs in B73 were defined as accessible (aUMR) or inaccessible (iUMR) based on its overlap with an accessible chromatin region in the B73 sample. For B73 aUMRs, the presence of an accessible region in the non-B73 genotypes was determined. The B73-based coordinates of the UMR in the corresponding genome were used to identify overlap with the ACRs defined in that genome. UMRs that overlap both an ACR in B73 and non-B73 genome were defined as stable ACRs. If the aUMR in B73 lacked accessibility in the non-B73 genome it was defined as B73-only ACR. Alternatively, if a UMR was inaccessible in B73 it could never be found accessible or show accessibility in the other genotype. If the iUMR lacked accessibility in the non-B73 genome, it was determined to have no ACR. If the sequence of the iUMR overlapped a defined ACR in the other genome, it was defined as a non-B73 ACR such that it was inaccessible in the B73 UMR but accessible in the shared sequence of Mo17, W22, or Oh43. The ACRs which were defined as either B73-only or nonB73-only were verified by assessing the 100bp cpm values within that region across the two genotypes.

## Data Availability Statement

Accessible chromatin data (ATAC-seq) generated for this study is available at NCBI short read archive under accession number PRJNA709664. In this study we also utilize previously published RNA-seq datasets that are available under accession numbers PRJNA657262 and PRJNA692023 and whole genome bisulfite datasets that are available under accession number PRJNA657677.

## Acknowledgments

This study was funded by support from the National Science Foundation IOS-1802848 to NMS and RJS, as well as IOS-1856627 to RJS. APM was supported by an NSF Postdoctoral Fellowship in Biology (DBI-1905869). We would like to acknowledge Christina Ethridge for preparation of ATAC-seq libraries and Pete Hermanson for preparation of WGBS libraries. The authors acknowledge the Minnesota Supercomputing Institute (MSI) at the University of Minnesota for providing computational resources that contributed to the research results reported within this paper.

---

**Figure S1:** A) Methylation state classification for each genotype based on alignment to their respective genome assemblies. Each 100bp bin of the genome was assigned a methylation state based on CG, CHG, and CHH methylation. Any bin with less than 2 cytosines was labeled “No Sites” and any bin with < 3x coverage was labeled “Missing Data”. For all other bins, context-specific cutoffs of methylation were used to classify CHH, CG only, CG/CHG, Intermediate and Unmethylated status. The proportion of each domain category for all bins in the respective genome are shown. B) The proportion of all bins for the B73 and non-B73 (Mo17, W22, Oh43) samples aligned to the B73v4 genome assembly that have coverage (black) or bins without enough coverage to be assessed (grey).

**Figure S2:** B73 ATAC-seq reproducibility. A) Metaplot of ATAC-seq coverage over annotated B73 genes for all ATAC-seq tissue samples aligned to the B73v4 genome assembly. The gene space was normalized to a 1kb region (represented in the middle of the metaplot) with the flanking upstream and downstream 1kb based on gene transcript direction. B) ATAC-seq was performed on two replicates for each genotype and ACR calls were generated for each sample individually (BN1A and BN2A) as well as the merged alignment file (BM merge). The venn diagram represents the overlap in defined ACRs for individual and merged samples for B73.

**Figure S3:** Overlap between ACRs and UMRs. The overlap between the Mo17 (A), W22 (B) and Oh43 (C) UMRs (blue) and ACRs (green) defined based on alignments to the B73v4 genome. Non-UMR ACRs that are defined as methylated are shown in parentheses below ACR count. RNAseq data for the same tissue sample was used to classify all B73 genes as not expressed (CPM < 1) or into expression quantiles of lowest expression (Q1) to highest expression (Q4). For each category of gene, the proportion of genes with aUMRs (D) and iUMRs (E) that are overlapping the gene or proximal to the gene(<2kb) was calculated.

**Figure S4:** Accessibility is often present only for a portion of the unmethylated region. (A-D) Several B73 UMRs are shown along with ATAC-seq data. IGV (Robinson et al., 2011) snapshots of the B73 genome showing ACRs within UMR space. Tracks include B73 gene and TE annotations, B73 methylation per cytosine in all contexts (CG: blue, CHG: red, CHH: yellow), B73 UMRs (black), B73 ACRs (blue), and B73 ACR coverage (grey).

